# Maximum Likelihood Localization Maps

**DOI:** 10.64898/2026.11.05.764521

**Authors:** Hossein Zarei Oshtolagh, Ronny Förster, Jonas Ries, Rainer Heintzmann

## Abstract

Single-molecule localization microscopy commonly reconstructs images from estimated emitter coordinates. We present maximum-likelihood localization maps (MLLM), a non-iterative approach that computes the positional likelihood of each detected molecule from the measured data and a supplied calibrated point-spread function, and adds it to an upsampled probability map, avoiding separate coordinate-based rendering. MLLM is fast, non-iterative, parallelizable, supports any calibrated point-spread function, compensates for local background and supports two- and three-dimensional reconstruction.

---

Single-molecule localization microscopy (SMLM) resolves cellular structures by combining observations of sparsely emitting fluorophores.^1,2^ Conventional workflows estimate individual emitter coordinates and subsequently render them into an image.^4^ Experimental PSF models have enabled localization beyond Gaussian approximations^3,5^, and iterative maximum-likelihood fitting supports efficient and precise coordinate estimation .^6,7^ Most PSF-fitting or non-iterative localization methods^9^ yield a single (most likely) coordinate per event. Probabilistic reconstruction approaches^10^ retain spatial uncertainty but can be computationally demanding. Deep-learning approaches avoid iterations at inference but require extensive training, and deliver either images^11^ or localizations,^12^, yet efficient reconstruction that directly retains each event’s probability-based positional likelihood is desirable. Here, we introduce maximum-likelihood localization maps (MLLM), a non-iterative approach that directly evaluates and accumulates spatial likelihood maps from measured SMLM data. MLLM produces reconstructed images without a separate coordinate-rendering step. MLLM builds on any measured or simulated PSF, accounting for or even using asymmetry, aberrations, and depth dependence. Since it does neither fit nor render events, it does not rely on a Gaussian approximation anywhere. The beauty of MLLM builds on the theoretical understanding of the information contained in a single detected photon as detailed below.

Like many other localization methods, we first identify and extract a region of interest (ROI) around each emitter and optionally determine its background value. As detailed in the Supplementary Information, the foundations of calculating the MLLM build on the insight that a point-mirrored PSF (*PSF’*) placed at the position of a single detected photon describes the spatial likelihood of a molecular emitter. A localization map is obtained by multiplying many such shifted molecular location *PSF’*s together and normalizing the result. If this procedure is described in log-likelihood rather than likelihood space, we obtain the log-likelihood result as a simple convolution of the detected image (in photon numbers) with the log-*PSF’*, which is both theoretically beautiful and fast to calculate. The final normalization can also be done in log space (see Supplementary Information for details), and a voxel-wise exponentiation (Figure 1) yields the single-event likelihood map to accumulate into the overall MLLM.

**Figure 1.**
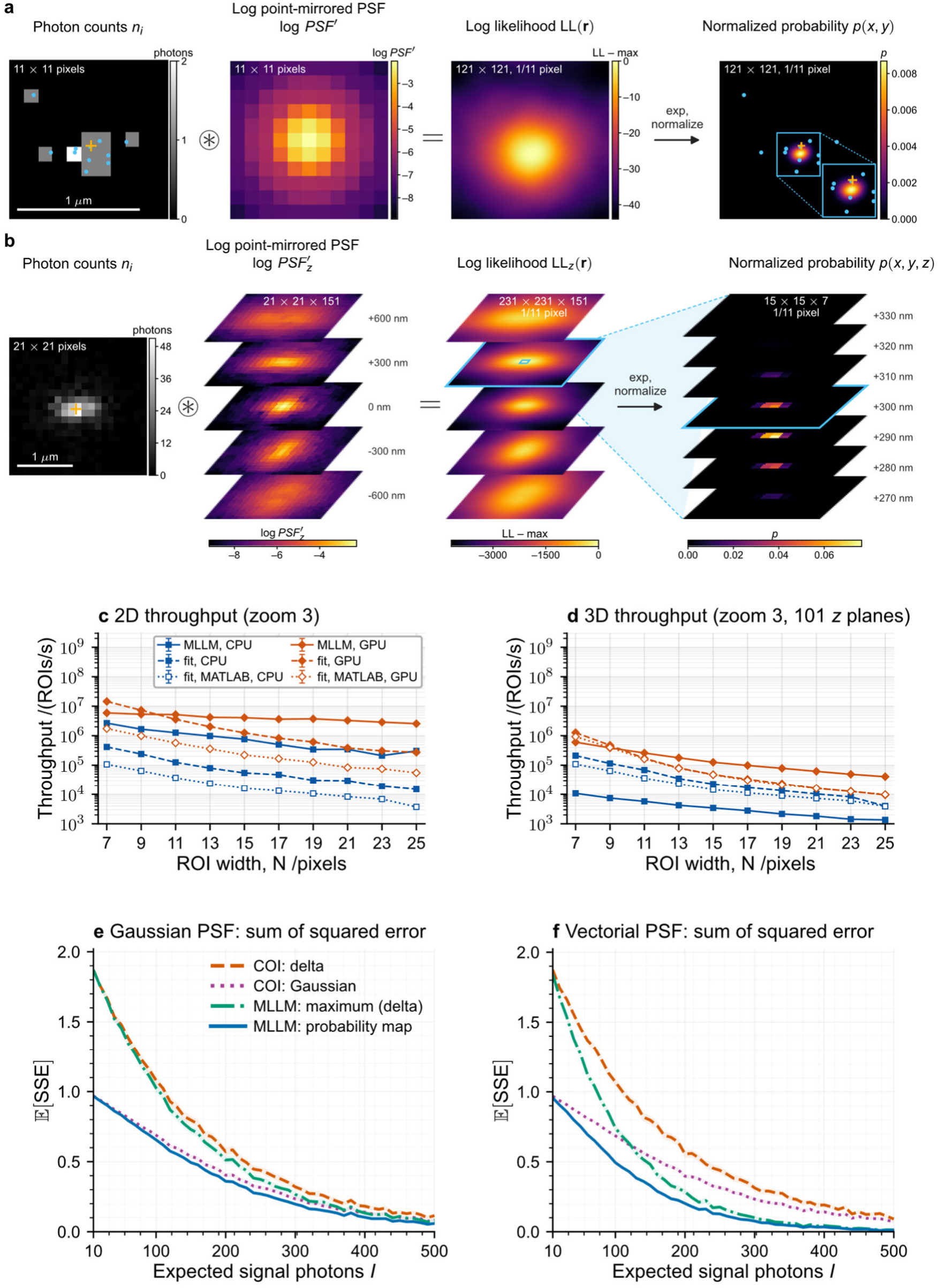
MLLM principle in 2D and 3D, throughput and map quality. a, 2D: a simulated in-focus emitter (vectorial PSF, 11×11-pixel ROI of 117 nm, 11 detected photons, no background). The photon counts are convolved (⊛) with the log point-mirrored PSF, log PSF′, giving the log likelihood LL; exponentiating and normalizing gives the probability map p(x, y). LL and p are 11× upsampled; the inset enlarges the framed 3×3-pixel region. Blue dots, detected photon positions; orange crosses, true position. b, 3D: the same process for an emitter at z = +300 nm (measured PSF, 21×21-pixel ROI, 951 detected photons) with log PSF′z of 151 planes 10 nm apart. p(x, y, z), normalized over the volume, is shown for the planes +270 to +330 nm around its maximum (blue guide; blue outline, true plane). Array sizes are given in each image. c,d, Throughput for 2D (c) and 3D (d; 101 planes) ROIs on a CPU (i7-11700KF, blue) and a GPU (RTX 3060, orange): MLLM (solid), Julia cSpline fitting (dashed; z fixed in c) and the original MATLAB/MEX cSpline software (dotted). 10,000 ROIs; medians and interquartile ranges of seven runs. e,f, Sum of squared error (SSE) of the maps for Gaussian (e) and vectorial (f) PSFs versus expected signal photons (4,000 ROIs per point; shading, 95% intervals). COI, center of intensity. The probability at the true bin is shown in Supplementary Figure S7. Scale bars, 1 3 µm.

To make the required substantial upsampling fast, we use an inverse chirp-z transform onto a finer lateral grid as the last step of the convolution calculation. A three-dimensional MLLM based on two-dimensional single-focus images is obtained via essentially the same algorithm, convolving a measured 2D image with each slice of a three-dimensional *PSF’* yielding a 3D result which is jointly normalized across lateral and axial candidate positions. A speed-optimized background compensation selects kernels from a precomputed bank based on the estimated signal-to-background ratio. Our implementation supports parallel computation on CPUs and GPUs. PSF normalization, finite support, periodic boundary conditions, and Fourier interpolation introduce assumptions and numerical approximations detailed in Online Methods.

To assess the benefit of retaining spatial likelihood information, we simulated 4,000 independent Poisson observations per signal level for Gaussian and high-NA vectorial PSFs, 10-500 expected signal photons, zero background, and an emitter fixed at the central bin (i.e. pixel) of a 31×31 grid of 25 nm candidate bins. We reconstructed each observation using the center of intensity (COI) rendered (the theoretically best unbiased estimator for purely Gaussian PSF and no background neglecting pixel integration) as either a delta function (histogramming) or Gaussian profile. Our fully normalized MLLM map was either compared directly or replaced by a delta at its maximum. We measured the mean reconstructed probability mass R at the true source bin index *c*, E[*R*_*c*_], and the sum of squared error (SSE), also called the Brier score^13^, E[*Q*],*Q* = ∑_*j*_(*R*_*j*_−*δ*_*jc*_)^2^with *δ*_*jc*_ = 1 for the true source bin and 0 otherwise (Figure 1e,f). Note the SSE, closely related to the root mean squared error (RMSE), is the appropriate metrics, since probabilities sum to one over the pixels.

A quantitative comparison for Gaussian and Vectorial PSF is shown in Figure 1 e,f where it is seen that MLLM outperforms classical position localization approaches in terms of SSE and E[*R*_*c*_] For a detailed discussion on why these norms are the appropriate way to mutually compare probability maps, i.e. rendered images with the ground truth, see Supplementary Material.

To benchmark computational efficiency, batches of 10,000 pre-extracted ROIs, timing Fourier transformation, likelihood evaluation, normalization, and reconstruction accumulation. In the MLLM-only campaign, 3D throughput on an NVIDIA RTX 3060 ranged from 3.90 × 10^4^ to 7.03 × 10^5^ ROIs/s across ROI widths of 7-25 pixels and zoom factors of 1-11, using 101 axial PSF planes. Timings excluded detection, preparation, and data transfers. Paired comparisons used a numerically validated Julia and MATLAB® (original code) implementation of iterative cSpline fitting^3^, including CRLB calculation and coordinate histogramming (Figure 1c, 2D; Figure 1d, 3D). These measurements compare workflows with different outputs.

To demonstrate reconstruction of experimental three-dimensional SMLM data, we applied MLLM to the publicly available SMLM challenge Tubulin-A647 dataset^8^. Processing 91,946 available frames produced a reconstruction from 3,572,756 accumulated detection maps (Figure 2). With reconstruction sampling of 10.64 nm×11.55 nm×10 nm, the resulting XY and XZ views reveal microtubule structures in three dimensions (Figure 2).

**Figure 2.**
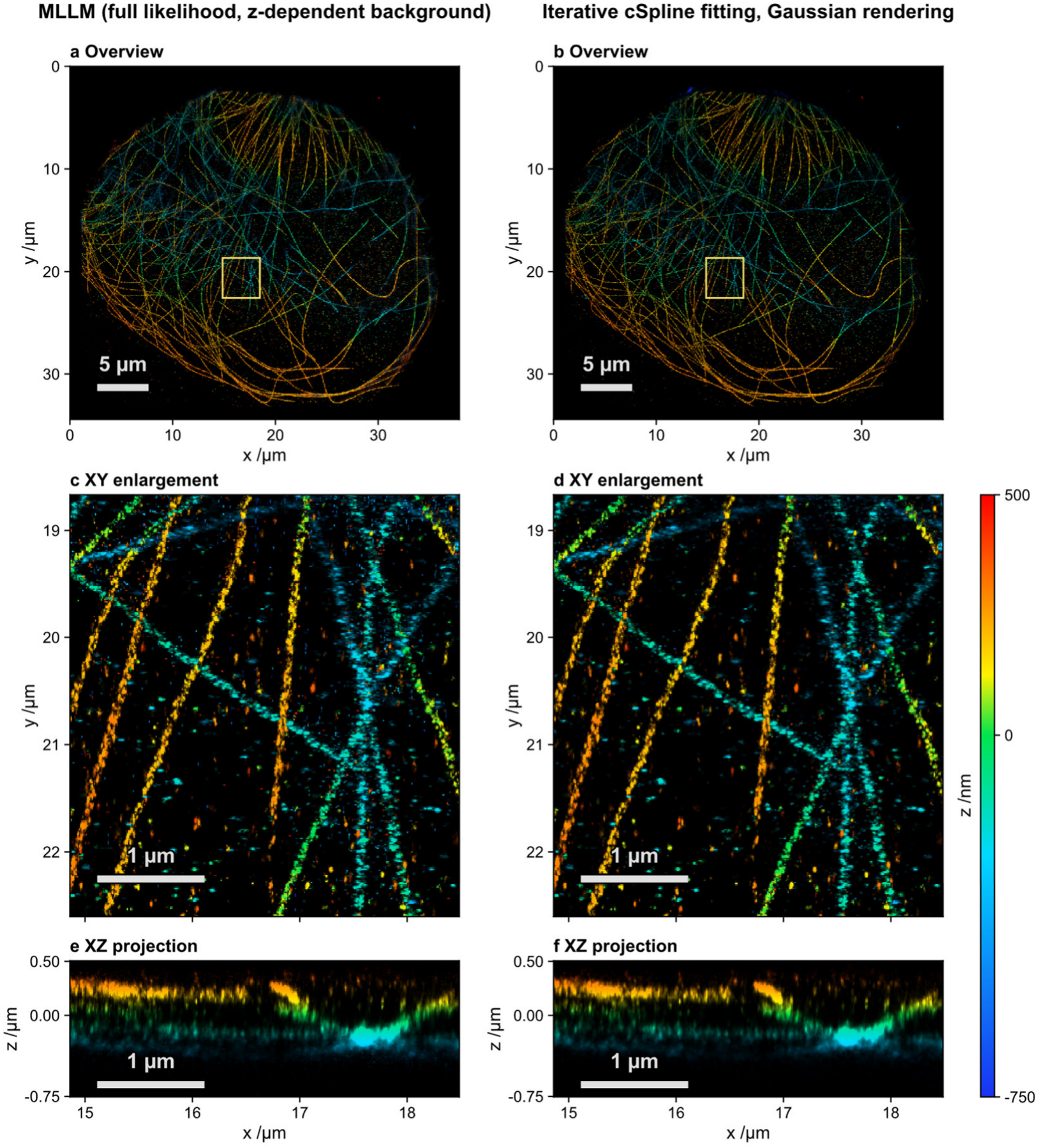
3D reconstruction of measured data with MLLM and iterative cSpline fitting. a,c,e, MLLM: each ROI adds its normalized 3D likelihood map. b,d,f, Iterative cSpline fitting (the authors’ MATLAB/GPU software^3^), each fit rendered as a 3D Gaussian with its CRLB widths. Both use the same 3,572,756 ROIs (21×21 pixels) of the SMLM Challenge Tubulin-A647-3D dataset^8^, the same calibrated PSF (Supplementary Figure S8) and the same lateral drift correction (Supplementary Figure S9). a,b, XY overviews. c,d, Enlargements of the boxed regions. e,f, XZ projections of the enlarged regions. Hue, depth (−750 to +500 nm). Scale bars, 5 µm (a,b) and 1 µm (c–f).

MLLM provides a direct framework for reconstructing spatial emitter-likelihood maps without iterative position fitting or successive separate coordinate rendering. Its implementation in FastMLLM.jl supports scalable two- and three-dimensional processing with calibrated PSFs. Quantitative accuracy depends on emitter isolation, detector sampling, PSF calibration, background estimation, and finite spatial support. MLLM replaced the ad-hoc Gaussian rendering of particle localizations into images with a theoretically founded probability map of molecular locations. This, along with the very fast non-iterative calculations, will advance applications of single-molecule localization microscopy.

## Online Methods

We aim to directly construct a probability map of possible emitter locations that fully accounts for Poisson statistics, and any known point spread function (PSF) without relying on simplified Gaussian-shape assumptions.

### Simplified case and assumptions

We consider a simplified case with a single emitter in a region of interest (ROI). This initial step replicates standard SMLM pipelines, where local intensity maxima are first identified in the recorded image, and a small ROI (or “patch”) around each such maximum is extracted. Each ROI is assumed to contain only one detected emitter, and potential multi-emitter regions are identified (e.g., by an upper threshold on the intensity integral) and discarded. To keep it simple, we first assumed zero background offset, out-of-focus light, and read noise. For this initial discussion, we also assume the acquired image is already sampled at the sampling of the final reconstructed probability map.

To explain the maximum likelihood localization map (MLLM) method, we first consider the case where only a single photon emitted from the emitter present in a ROI was detected. We then model the emitter’s position probability distribution by placing the PSF, point-mirrored at the nominal emitter position (i.e., where the photon was detected). The point-mirrored PSF appears, since we need to account for potential emitter positions having caused our detected photon. This matters for non-centro-symmetric shapes. Thinking of a shifted PSF, makes it easy to see that estimating possible emitter positions requires reversing this shift, which is exactly what point-mirroring achieves. Mathematically, this is the adjoint convolution with the PSF operator. We further assume that the PSF is normalized to its integral, so that the result directly corresponds to a spatial probability density function (PDF) of the detected photon.

Let us now consider a second photon, detected at another location on the sensor. We repeat the same operation, yielding a second probability map. To combine our acquired information, having detected photon one and photon two, we now need to multiply these two maps elementwise and normalize the result to reflect the overall emitter position probability distribution given the two detected photons.

Generalizing this concept to many detected photons and across the full ROI, we treat each photon detection event as an independent contribution to the emitter probability distribution. In a practical array detector, a given pixel *i,j* recorded *N(i, j)* photons. This is equivalent to observing a photon at this location N independent times.

Therefore, the overall emitter probability distribution can be modeled as the product of point-mirrored PSFs, each centered at the corresponding detection site and raised to the power of the number of detected photons in that pixel. This leads to the following expression for the probability map:

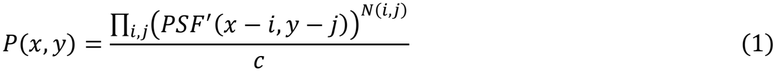

Here, *N(i, j)*denotes the photon count recorded at pixel *(i, j), PSF’(x − i, y − j)* represents the PSF flipped and its origin shifted to that location, and *c* is the normalizing constant. This formulation effectively collects spatial information from all detected photons, assuming they originate from a single emitter. Up to a position-independent multinomial factor, the numerator is the likelihood of the observed photon counts for an emitter at *(x, y)*. To obtain a valid probability map, *c* is chosen such that its integral amounts to one, *c = ∬ ∏*_*ij*_*PSF′(x − i, y − j)*^*N*^*(i, j)dxdy*. Assuming a uniform prior over candidate emitter positions, *c* is therefore proportional to the marginal likelihood (evidence) of the data, and *P(x, y)* is the posterior probability distribution of the emitter position. We then have

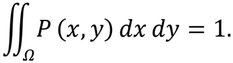

Directly evaluating this product over all image pixels can be computationally intensive and numerically unstable, particularly at high photon counts and spatially wide PSFs. As typically done in the ML framework, we therefore apply the natural logarithm to the unnormalized probability expression. This transforms the product into a sum. Taking the logarithm on both sides yields the following expression:

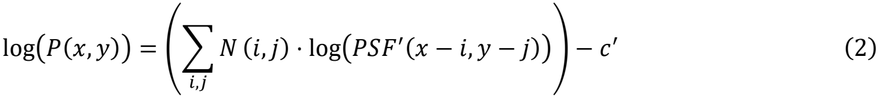

with *c*^’^ = lo*g*(*c*).

This sum is a convolution between the recorded image (in photon number) and the logarithm of the point-mirrored PSF.

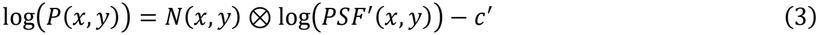

According to the convolution theorem, this operation can be computed efficiently in the Fourier domain:

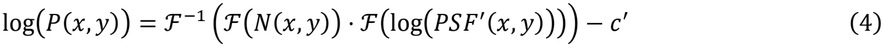

where ℱ ( ) represents the Fourier transform.

Finally, the emitter probability map is calculated by applying the exponential function elementwise to the result:

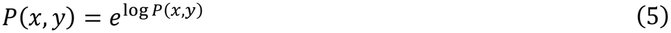

This procedure (closely related to the “soft-max” operation in neural networks) typically yields a smooth estimate of the emitter’s possible locations based solely on a single convolution operation, while maintaining the physical correctness of the photon statistics and avoiding any assumption about the PSF being Gaussian-shaped, with the detected intensity image *I(x, y)*, measured in ADU and the gain factor g, or even having a finite variance. As detailed in the supplement, the gain factor g potentiates the result probability and therefore needs to be accounted for accurately when converting the ADUs to effective photons:

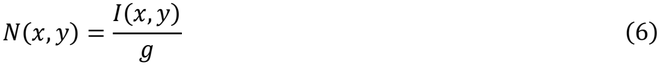

If we replace the soft-max operation above with a “one-hot” operation, placing a one only at the location of the maximum pixel, we essentially obtain a non-iterative maximum likelihood fit binned (histogrammed) onto our final reconstructed grid.

### Upsampling to the reconstruction grid

Raw SMLM data are acquired on a detector grid that is typically much coarser than the desired sampling of the reconstructed localization map. To obtain a practically useful MLLM implementation, we evaluated the likelihood on a user-defined finer spatial grid while still retaining the point-spread function (PSF) on the original camera grid.

For sinc upsampling during processing, we first transformed the camera-grid ROI to Fourier space using the Fast Fourier Transform (FFT) and multiplied by the Fourier-domain kernel, i.e., the FFT of the logarithm of the point-mirrored PSF. Instead of explicitly zero-padding this Fourier-domain array and applying an inverse Fourier transform on a larger grid, we directly evaluate the result at the desired fine-grid coordinates using a centered inverse chirp z-transform (iCZT)^14^. This combines convolution and fine-grid evaluation without explicitly upsampling the ROI or the PSF.

To assess sensitivity to the sampling approximation, we also evaluated uniform and sparse count mappings in two-dimensional, zero-background controls. Uniform mapping divides each recorded detector-pixel count equally among its associated fine-grid subpixels, whereas sparse mapping assigns the full count to the detector-pixel center. These mappings define precomputed linear operators that act directly on the camera-grid ROI, avoiding explicit construction of an upsampled image. They differ from physical detector integration: uniform mapping averages log-PSF values over a pixel footprint, whereas the detector-level Poisson likelihood requires the logarithm of the pixel-integrated predicted intensity. Sparse mapping samples the interpolated camera-grid PSF at detector-pixel centers and approximates this likelihood, provided the supplied PSF incorporates pixel integration and accurately represents fractional shifts. We therefore used these mappings to numerically compare independently evaluated detector-level likelihoods, rather than treating uniform redistribution as exact pixel integration (Supplementary Figure S6). The principal reconstruction and throughput results use the *sinc*/iCZT pathway, which interpolates the camera-grid positional score.

### Implementation

We first converted the raw data into effective photon numbers using the known camera gain and offset. These parameters can be obtained from camera calibration data or estimated from recorded images^15^. We subtracted the offset and divided the resulting intensities by the gain before further processing.

For end-to-end reconstruction, we applied a Difference-of-Gaussians (DoG) filter to each camera-corrected frame and detected local maxima at integer camera-pixel positions. The filter suppressed slowly varying background contributions, with widths chosen based on the approximate PSF size. We extracted centered ROIs from the corrected, unfiltered frames and retained their integer anchors and frame indices for subsequent processing. This ROI-based implementation supported event-specific background-kernel selection and independent normalization of each detection’s likelihood map before accumulation. The single-emitter model assumed that neighboring emitters contributed negligibly within each ROI.

The main reconstruction assumed that each extracted ROI contained a single active emitter. We also implemented a two-emitter likelihood formulation for overlapping emitters (see Supplementary Information). For the reconstructions presented here, we used the computationally more efficient single-emitter ROI-based approach.

For each ROI, we calculated a two-dimensional FFT and multiplied the resulting spectrum by the Fourier-domain log-PSF kernel. For *sinc*-kernel resampling onto the reconstruction grid spacing, we evaluated the resulting Fourier-domain log-likelihood directly on the fine reconstruction grid using the centered iCZT described below. We converted the log-likelihood map into a normalized localization likelihood in a numerically stable manner by first subtracting its maximum value, applying the exponential function, and then normalizing the resulting map to unit sum. We then accumulated the normalized local likelihood maps at their corresponding positions in the final reconstruction.

### Background compensation

To account for nonzero background, we used a precomputed bank of likelihood kernels dependent on the background-to-signal ratio, with the background estimated separately for each axial hypothesis. For each axial plane z of the PSF model, normalized to unit sum within the ROI, we precomputed the fraction of PSF intensity falling on the one-pixel-wide outer border, *β*_*z*_ = ∑_*i* ∈∂RO*I*_*p*_*z*_ (*i*), counting each corner once. For each camera-corrected, unfiltered *N*× *N* ROI with corrected pixel counts, we calculated the total and border counts

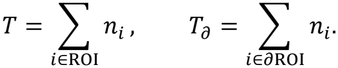

Assuming an emitter at depth *z* with signal *S* and a uniform background b per pixel, their expected values are *T* = *S* + *N*^2^*b* and *T*_∂_ = *β*_*z*_*S* + (4*N*− 4)*b*. Solving these two equations gave, for each axial plane,

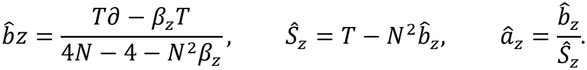

For every axial plane, *â*_*z*_ selected the nearest entry in a precomputed bank of kernels lo*g* (*P*_*z*_ + *A*)/(1 + *N*^2^ *A*), logarithmically spaced in *a*. We assigned ratios outside the bank’s range to the corresponding endpoint. Different axial planes of the same ROI could therefore use different background levels; the normalization term keeps their likelihoods comparable.

To reduce noise at low photon counts, *â*_*z*_ was refined using all ROI pixels. With *P* = *N*^2^ and the signal fraction w, the model predicts the fraction *q*_*i*_ = *w p*_*z*_(*i*) + (1 − *w*)/*p* of the total count in pixel *i*, assuming an emitter at the ROI center. Starting from *w*_O_ = *Ŝz*/*T*, we applied one Newton (Fisher-scoring) step,

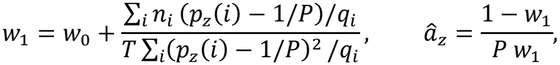

where *q*_*i*_ is evaluated at *w*_0_, and *w*_0_ and *w*_1_ are restricted to [0.02, 0.98] and [10−3, 1], respectively. Starting from a consistent estimate, a single step is asymptotically as efficient as the maximum-likelihood estimate; further iterations did not change the results.

The estimated background was used for kernel selection rather than subtracted from the ROI counts entering likelihood evaluation. This procedure required no ground-truth signal or background values and did not jointly fit these parameters. Unlike a border mean, it does not attribute defocused PSF tails on the border to background. It assumed a spatially uniform background and a single emitter per ROI. Our implementation provides single-threaded CPU, multithreaded CPU, and CUDA backends. We used the same mathematical likelihood model for all backends.

### Three-dimensional reconstruction

We extended the MLLM formulation to three-dimensional localization using a measured or simulated three-dimensional PSF stack. Each detected emitter still had a two-dimensional ROI as input, while the PSF stack represents the expected emitter image at a series of axial positions.

For the experimental Tubulin-A647-3D reconstruction, we used the fluorescent-bead calibration stack supplied with the SMLM Challenge^8^. Individual bead images were background-subtracted, aligned with subpixel precision, and averaged. To reduce residual calibration noise, we retrieved the pupil function from the averaged PSF using a difference-map phase-retrieval algorithm^16^. We then forward-propagated the recovered pupil to generate a smooth model PSF stack. This procedure limited the influence of calibration noise on comparisons between axial hypotheses. The recovered pupil phase is shown in Supplementary Figure S8a.

For each axial plane *z*, we used a normalized PSF kernel *q*_*z*_, including the background contribution described in the Background compensation subsection. At zero background, *q*_*z*_ reduces to the normalized PSF. With *N*_*ij*_ denoting the measured photon counts and *q*′_*z*_(*x, y*) = *q*_*z*_(−*x*, −*y*) the point-mirrored kernel, the positional log-likelihood score on the camera grid was

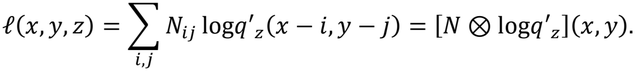

We computed this lateral convolution for each axial hypothesis by multiplying the Fourier-transformed ROI by the corresponding Fourier-transformed log-kernel. The resulting score was evaluated on the finer lateral reconstruction grid using the centered inverse chirp-z transform, while the axial sampling followed the spacing of the supplied PSF stack.

Finally, we exponentiated and normalized the scores jointly over all lateral and axial candidate positions:

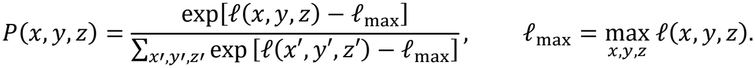

Subtracting the common maximum ensured numerical stability. Joint normalization preserved the relative likelihoods across axial planes and yielded a three-dimensional localization map for each detected event.

This formulation did not require fitting separate PSF parameters such as the widths of an astigmatic emitter image. Instead, the complete calibrated PSF shape at each axial position entered directly into the likelihood calculation. Axial information encoded in the PSF, for example through astigmatism or other depth-dependent PSF variations, was therefore incorporated directly through the supplied PSF stack.

### Centered inverse CZT implementation

We evaluated inverse spatial Fourier transforms using centered chirp z-transforms implemented on both CPU and GPU. For an *N×N* Fourier-domain ROI and an output grid of size *M×M*, the one-dimensional transform matrix was defined as

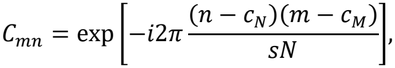

where s is the spatial zoom factor and *c*_*N*_and *c*_*M*_ denote the center indices of the source and destination grids, respectively. The two-dimensional transformation was evaluated separably as

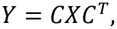

where X denotes the Fourier-domain ROI.

The corresponding inverse transform was obtained using the conjugation identity

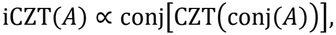

with the appropriate normalization for the discrete transform convention.

## Acknowledgements

R.H. and H.Z. were supported in this work by the DFG-funded Collaborative Research Center PolyTarget (SFB 1278, Project ID: 316213987), Project Area C, Project C04. H.Z. thanks Rahel Koch for helpful discussions and constructive suggestions about this work.

## Data availability

The experimental 3D data (Tubulin-A647-3D, including the bead calibration stack) are from the 2016 SMLM Challenge and are publicly available from the challenge website (https://srm.epfl.ch)^8^. The experimental 2D dSTORM data of microtubules in U2OS cells were published by Diekmann et al.^17^. All simulated data were generated with the code described under Code availability, using the parameters given in the Online Methods.

## Code availability

FastMLLM.jl, the Julia implementation of MLLM (CPU and CUDA GPU), is available at https://github.com/hzarei4/FastMLLM.jl under the MIT license. The version used in this study, including the scripts that generate all figures, simulations and throughput benchmarks, will be deposited at Zenodo upon publication. The iterative cSpline comparisons used the authors’ original software (SMAP/cSpline, Li et al.^3^) https://github.com/jries/fit3Dcspline and a Julia reimplementation included in the repository. ThunderSTORM^18^ (ImageJ^19^) was used for the 2D comparison.

## Supplementary materials

### Scaling the photon number (camera gain)

In practical imaging systems, the absolute detected photon count *N*(*x, y*) depending on the pixel coordinates *x, y* is often scaled by a gain factor *g*, in analog-to-digital units (ADU), such that the observed intensity *I*(*x, y*) at each pixel is given by

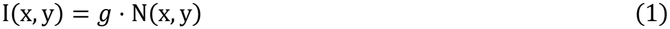

Inserting eqn. 6 in the main text into eqn. 3 in the main text, we obtain

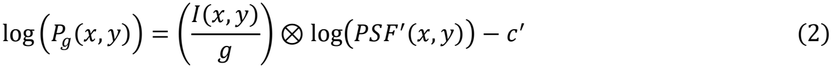

Since *g* is a multiplicative scalar coefficient and the convolution operation is linear, we obtain

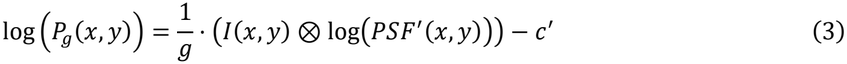

Now, considering the scaling effect of *g*, we obtain the probability expression

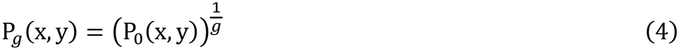

At unity scaling, we obtain

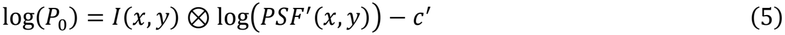

Since *g* affects the exponent of the result probability, different values of *g* affect the resultant map significantly. Since the gain factor scales the exponent of the likelihood, an incorrect gain changes the width and normalized peak value of the reconstructed distribution. For an underestimated photon number, the distribution becomes broader, whereas an overestimated photon number produces an artificially narrower distribution. In the idealized case, the position of the likelihood maximum is unaffected by positive global scaling, but the shape of the normalized likelihood map depends directly on the gain calibration.

### Extension to overlapping emitters

As an extension to the work presented in the main text, we here consider localizing two closely spaced, simultaneously emitting molecules whose PSFs overlap within the same ROI. The principle underlying MLLM can distinguish between the two, even when their signals overlap.

#### Two emitters in 1D

We start with a one-dimensional problem with two emitters. We simulated two emitters in one dimension using a full-vectorial, high-NA PSF (λ = 500 nm, NA 1.4, refractive index 1.52). We set the sampling to 20 nm per pixel and placed the emitters at −80 nm and +80 nm relative to the center of the frame. The distance between the two emitters was therefore 160 nm, below the Sparrow limit of 168 nm for this PSF. Each emitter had an expected number of 2,000 detected photons, and we added a background flux of 5 photons per pixel. After applying Poisson noise, the total number of detected photons in this simulation was 4,336.

In analogy to the arguments made for the single molecule localization, we now consider what it means to detect a photon under the assumption of two overlapping emitters. Since we do not know from which emitter a detected photon originated, we assume that it originated from either of the two with the same likelihood. The role of the PSF, here a 1D function is now taken by a two-dimensional joint probability map. Since a detection refers to an “or” operation, we construct this two emitters’ joint probability map as a normalized sum of the individual PSFs. In this two-dimensional joint probability map (Figure S1), each dimension represents the position of a respective emitter *x*_*i*_.

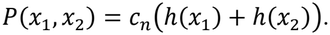

Where *h*(*x*) refers to the particle’s localization probability density function (i.e. the flipped PSF’), assuming an emitted centered at the origin and *c*_*n*_ is a normalization constant.

**Figure S1.**
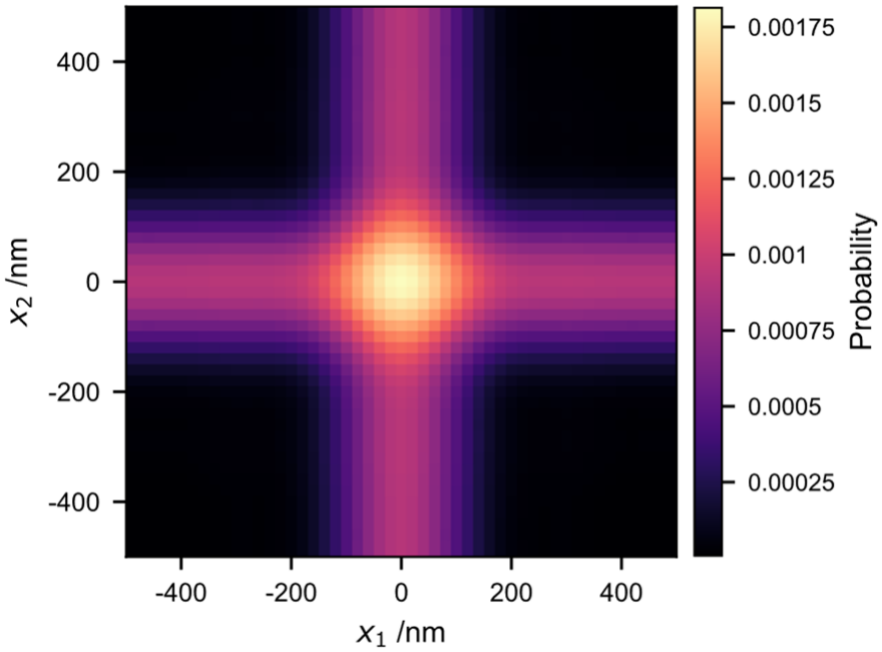
Joint-probability map of the two particles in 1D

We detect a single photon in this space; the position of this single photon in joint probability space will be at the diagonal where *x*_*i*_ = *x*_*j*_, since we do not know which particle it belonged to. Like in the single-emitter case we now also need to convolve this delta peak with the joint probability PSF’ as given in Figure S1. Detecting a second photon, we also multiply this joint probability map by a shifted one, centered at the detected photon’s place on the diagonal. We repeated this for all the detected photons from two emitters. In log space, this multiplication process is again a convolution.

To process the measured data, we projected it to this two-dimensional space by mapping to the diagonal. So, each element on the diagonal axis is *x*_*i*_ = *x*_*j*_and the value is the detected photon number.

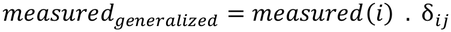

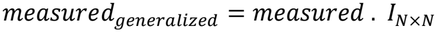

Where *I* is the identity matrix of size N by N, where N is the measurement array size.

Like in the one-emitter case (see main text of this paper), we calculated the convolution of this generalized measurement with the logarithm of the joint probability map of two emitters in 1D. To get each particle’s likelihood map, we calculate the marginal probability by summing over the dimensions *x*_*i*_ and *x*_*j*_.

Due to ambiguity in the particle order, the result matrix is symmetric along the diagonal. As a result, the marginal probabilities of the two particles are identical, and this degeneracy remains plausible.

After incorporating the detected photons, we obtained a normalized joint likelihood *P*(*x*_1_, *x*_2_)for the positions of the two emitters. Because the emitters are indistinguishable, the joint likelihood is symmetric under exchange of the emitter coordinates,

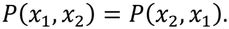

To obtain a one-dimensional localization map, we marginalized the joint likelihood over either emitter coordinate. The marginal likelihoods were calculated as

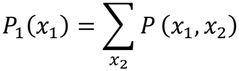

and

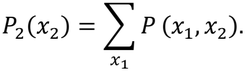

Owing to the exchange symmetry of the two identical emitters, these two marginal distributions are identical, and each contains the two possible emitter positions.

Figure S2 shows a simulation of photon detection from overlapping emitters with the parameters detailed above. The Sparrow limit distance for this PSF is 168 nm (while the Rayleigh limit is 218 nm), and the emitters are placed 160 nm apart. It is seen that they can still be localized if processed accordingly in joint-probability space.

**Figure S2.**
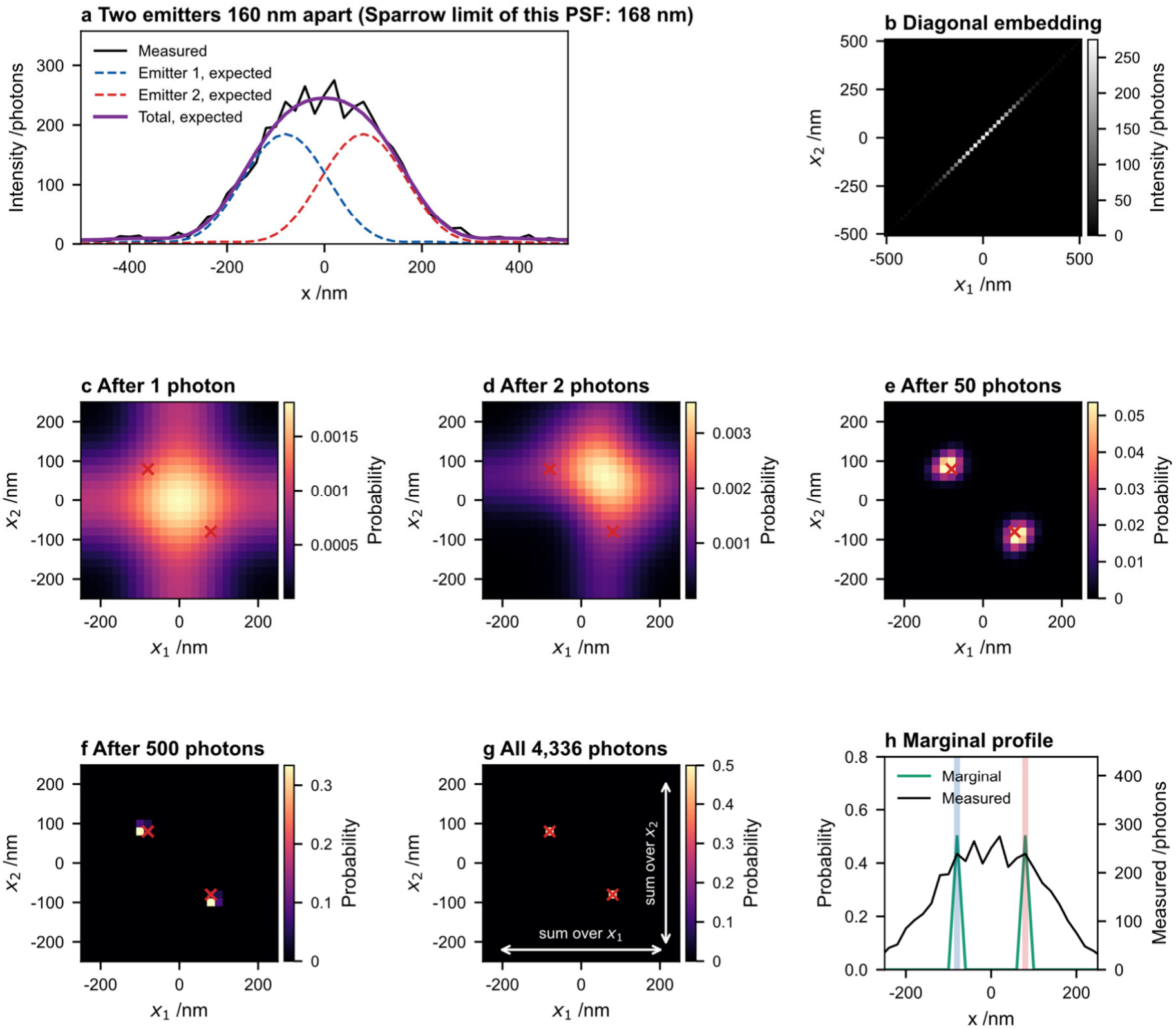
Joint-likelihood construction and marginalization for two overlapping emitters in one dimension. a, Simulated probability distributions of two emitters separated by 160 nm and the corresponding noisy (Poisson-distributed) measured intensity profile; b, the measured one-dimensional photon-count distribution embedded along the diagonal of the two-dimensional (*x*_1_, *x*_2_) coordinate space. c–f, Evolution of the joint two-emitter likelihood as the detected photon information is incorporated, progressively concentrating the likelihood around the two exchange-symmetric emitter configurations. Red crosses indicate the true emitter-coordinate combinations; g, final joint likelihood and directions of marginalization over either emitter coordinate; h, resulting one-dimensional marginal localization profile, shown together with the measured intensity distribution and the true emitter positions. Because the two emitters are indistinguishable, the joint likelihood is symmetric under *x*_2_ ↔ *x*_3_, and the two marginal likelihoods are equivalent.

#### Two emitters in 2D

We extended the same construction to two emitters in a two-dimensional image. For two equal-brightness emitters at positions ***r***_1_ = (*x*_2_, *y*_3_) and ***r***_2_ = (*x*_1_, *y*_2_), the probability of detecting a photon at position *q* is proportional to

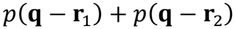

where *p* denotes the PSF. The corresponding joint localization likelihood therefore depends on four spatial coordinates,

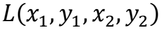

To evaluate this likelihood using the same convolution formulation as for the single-emitter case, we embedded the measured two-dimensional photon-count image *I*(*x, y*)into a four-dimensional diagonal tensor,

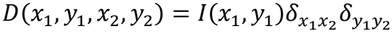

where denotes the Kronecker delta. Thus, the measured *N*× *N*image occupied only the diagonal subspace *x*_1_ = *x*_2_ and *y*_1_ = *y*_2_ of the *N*× *N*× *N*× *N*tensor.

We then evaluated the four-dimensional log-likelihood by convolving this embedded measurement with the logarithm of the two-emitter joint kernel,

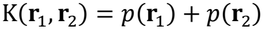

After exponentiation and normalization over the complete four-dimensional space, we obtained the joint probability distribution of the two emitter positions. Because the two emitters are indistinguishable, the joint distribution is invariant under exchange of the two emitter coordinates,

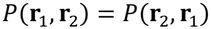

We therefore reduced the four-dimensional distribution to a two-dimensional localization map by marginalizing over the coordinates of one emitter,

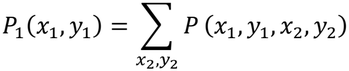

The corresponding marginal for the second emitter is identical because of the exchange symmetry. Consequently, the two peaks in the marginal map represent the two possible emitter positions without assigning an arbitrary label to either molecule.

For the example shown in Figure S3, we simulated two emitters with a Gaussian PSF (σ = 0.51λ/NA/2.355 = 77.3 nm for λ = 500 nm and NA 1.4, integrated over each camera pixel), 50 nm per camera pixel, an expected signal of 100 photons per emitter and zero background; after Poisson noise, the image contained 207 detected photons. The emitters were placed 166.7 nm apart, just below the Sparrow limit of this PSF (0.47λ/NA = 167.9 nm; Rayleigh limit 0.61λ/NA = 217.9 nm). Candidate positions were evaluated on a 16.7 nm grid (one third of a camera pixel). The reconstructed marginal likelihood recovered two distinct probability maxima, each within one grid bin of the corresponding simulated emitter position.

**Figure S3.**
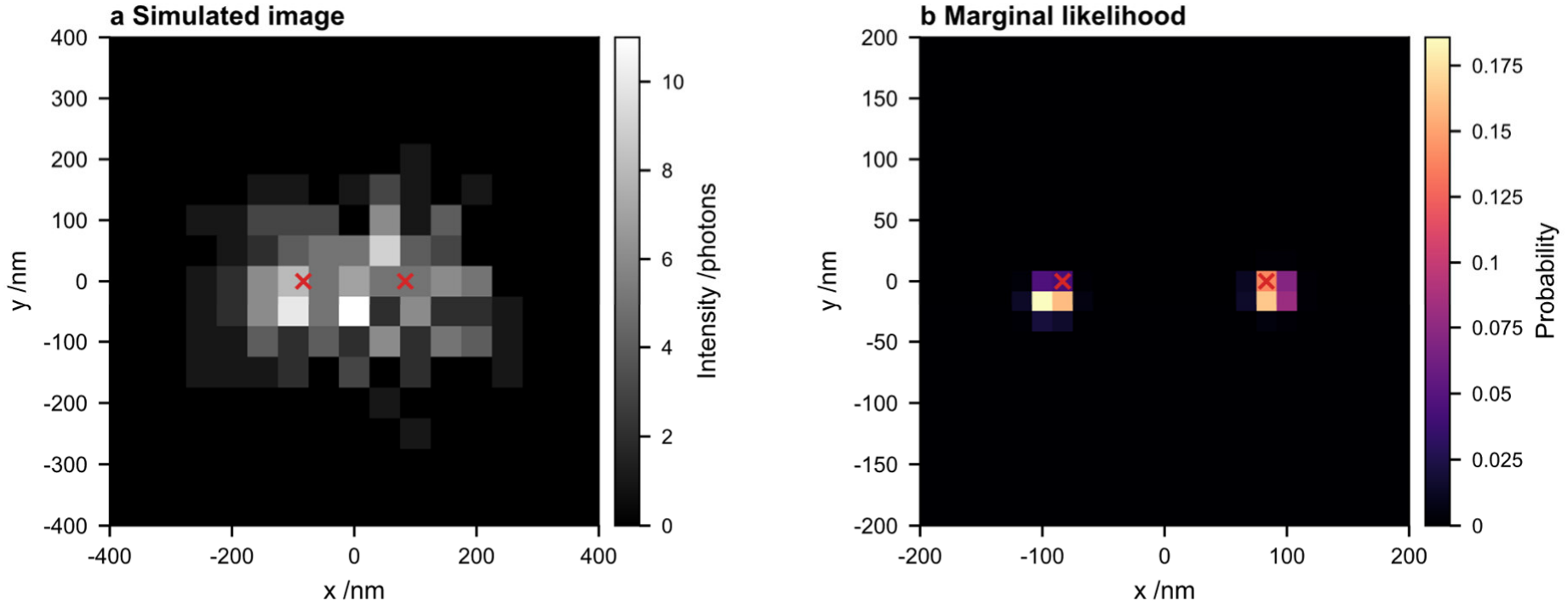
Two-emitter MLLM reconstruction in two dimensions at the Sparrow limit. a, Simulated image of two emitters separated by 166.7 nm (Sparrow limit 167.9 nm), with an expected signal of 100 photons per emitter, zero background and a camera sampling of 50 nm per pixel (207 detected photons). b, Marginal localization likelihood obtained from the four-dimensional joint two-emitter likelihood after summation over the coordinates of one emitter, on a 16.7-nm grid. Owing to exchange symmetry between the two indistinguishable emitters, the marginal distribution contains both possible emitter positions. Crosses, true positions.

Note also that we assumed the emitters to be equally bright. This assumption can be violated in practical situations (e.g. when an emitter bleaches during the acquisition of a frame), with potentially strong effects on the resulting probability maps. Although we can see that the MLLM concept can be extended to multiple overlapping emitters, this effect and the algorithmic slow down by the higher dimensionality coined us to not include localization of multiple overlapping emitters in the data evaluation at the current state.

### 2D reconstruction of a measured dataset

To validate the practical applicability of the MLLM algorithm, we applied it to a publicly available dataset from Diekmann et al.^17^. From this dataset, we used a sequence that contains wide-field dSTORM data of fixed U2OS microtubules immuno-stained with Alexa Fluor 647 and imaged in a glucose-oxidase/catalase buffer supplemented with 143 mM *β*-mercaptoethanol. The 642 nm excitation beam was adjusted to 46 mW at the objective (roughly 6.4 kW cm^−2^) while the camera integration time was kept at 30 ms.

A 405 nm activation line, limited to 20 % of its 200-mW maximum, was applied intermittently to sustain a constant density of switchable fluorophores throughout the 31,000-frame acquisition. We discarded the first 1000 frames since these frames were too crowded but forced most of the fluorophores to the dark state. Then we scaled this data to the photon numbers using the algorithm described in Heintzmann et.al^15^.

These measurements show a median integrated signal of about 3,000 photons per localization (interquartile range 1,400–5,400 photons; ThunderSTORM fits converted with the calibrated gain of 0.516 photons/ADU), so each event carries a comfortable signal-to-noise ratio for high-precision localization.

**Figure S4.**
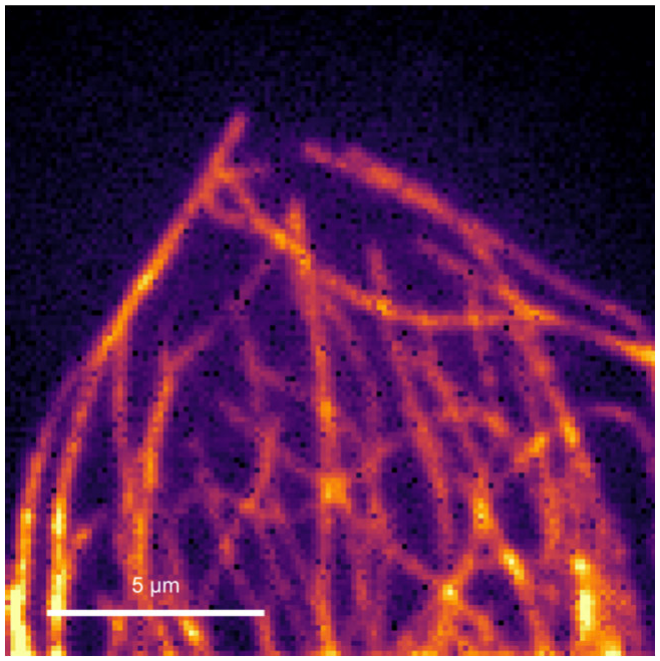
The sum image (widefield image) of all 30,000 frames of the measured data. The background offset was subtracted, and contrast was enhanced. Scale bar is 5 μm.

We compare our reconstructed scene with the reconstruction of the ThunderSTORM^18^ plugin for ImageJ^19^. Both reconstructions were performed assuming a gain of 0.516 photons/ADU, estimated based on the single frame image gain estimation^15^ averaged over all frames accounted for in the reconstruction. The offset was estimated to be 96.6 ADU. The comparison of the reconstructions is shown in Figure S5.

**Figure S5.**
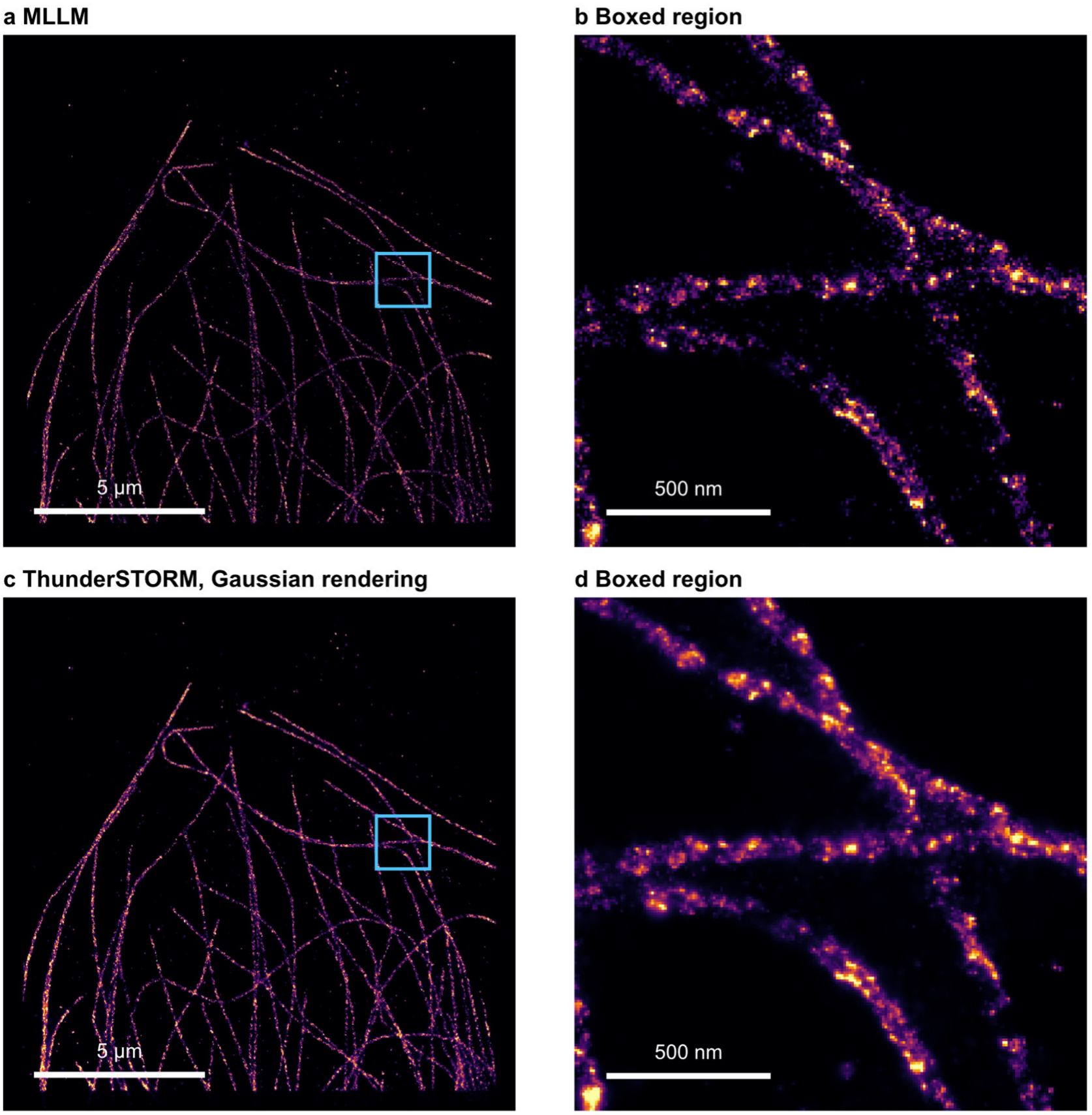
Comparison of two reconstructions of the dataset. a) MLLM reconstruction result b) a 200 by 200 pixels cropped region of the MLLM reconstruction c) ThunderSTORM’s reconstruction with normalized Gaussian rendering d) a 200 by 200 pixels cropped region of the ThunderSTORM’s reconstruction with normalized Gaussian rendering. The small regions are indicated with a red square. Scale bars on the left are 5 μm, and on the right are 500 nm

As it is depicted in Figure S4 (Widefield sum) and the reconstructions in Figure S5, the two sides of tubular structure of the tubulins are visible in both reconstructions. The number of events detected in the MLLM reconstruction was 477,495, and for the ThunderSTORM reconstruction was 817,464.

### Localization statistics

In addition to the full vectorial PSF, we also approximated the system PSF by a circularly symmetric Gaussian whose width we approximated by a low NA equation obtained from the Airy pattern^20^, depending solely on the emission wavelength *λ* and the numerical aperture (*NA*):,

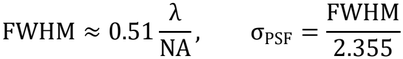

This approximate value of *σ*_*PSF*_ *w*as used for the Gaussian-shaped PSF in the simulations.

To determine the reconstruction quality of the MLLM algorithm, we ran 1,000 trials for each simulation condition. In every trial, we generated Poisson photon counts with an expected total photon number (100 and 400), reconstructed the MLLM probability map, and calculated the sum of squared errors (SSE). The comparison is shown in Figure S6.

**Figure S6.**
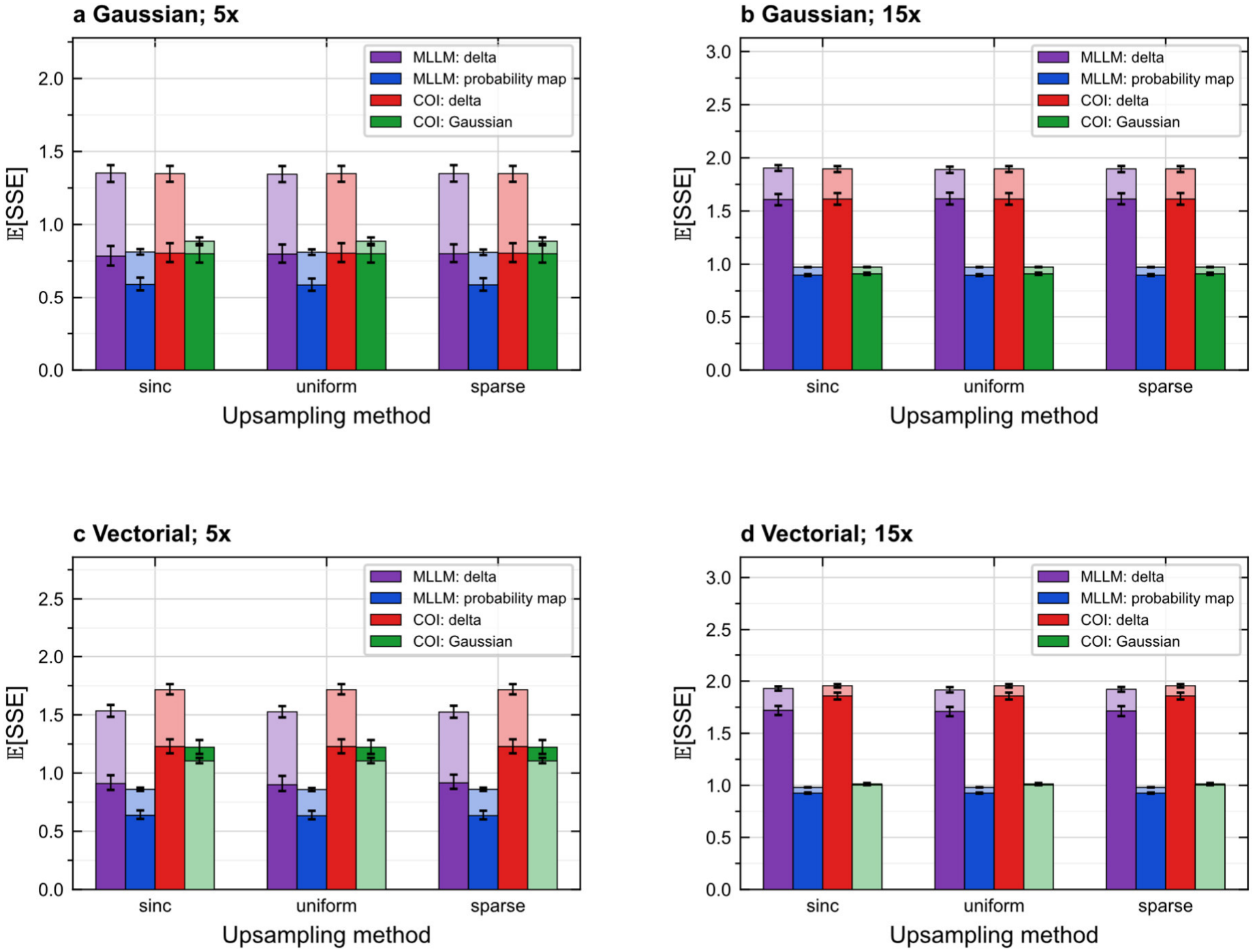
Sum of squared error of the maps in the randomized-position study. We simulated 1,000 ROIs per condition (17×17 pixels of 100 nm, emitter at a random sub-pixel position, no background), reconstructed them with the sinc, uniform and sparse upsampling methods, and scored each map over all bins by SSE = Σ_*j*_ (R_*j*_ − δ_*j*_c)^2^ (R, normalized map; c, true source bin; lower is better). a,b, Gaussian PSF; c,d, vectorial PSF; upsampling factor 5 (20-nm bins) and 15 (6.7-nm bins). Bars show the MLLM maximum as a delta (purple), the MLLM probability map (blue), the COI as a delta (red) and the COI Gaussian rendering (green); full color, 400 photons; faint, 100 photons. Error bars, bootstrap 95% intervals. The MLLM map has the lowest SSE in every condition.

### Mean true-bin probability

We did not consider the Kullback-Leibler divergence for comparing the distributions, since the histogrammed renderings would lead to frequent infinities (e.g. if the ground truth is zero but the map is non-zero in a pixel). At every photon count, retaining the full MLLM map gave the lowest SSE of all four methods, even though it assigned less mass to the true bin than maximum-only reconstruction. At 10 expected photons, the mean SSE decreased from 1.869 (maximum only) to 0.967 (full map) for the Gaussian PSF and from 1.841 to 0.957 for the vectorial PSF (Figure 1 e,f). Compared with the Gaussian-rendered COI, the reduction was largest at intermediate photon counts. At 100 expected photons it was from 0.689 to 0.656 for the Gaussian PSF and from 0.686 to 0.494 for the vectorial PSF. As photon counts increased, true-bin mass approached unity and SSE approached zero. These results demonstrate a benefit of retaining positional uncertainty in this photon-limited, fixed-center experiment.

**Figure S7.**
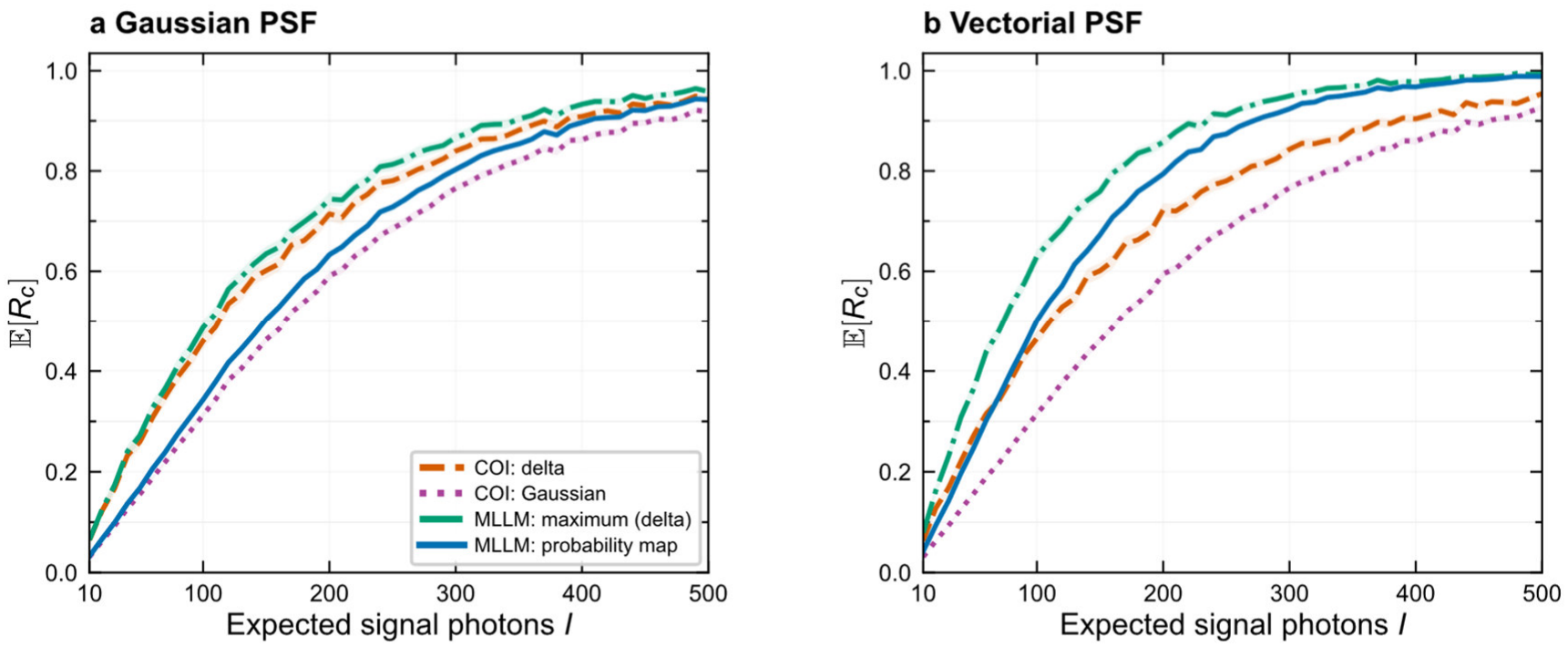
Probability at the true bin. Expected probability in the true bin, *E*[*R*_*c*_], versus expected signal photons for a Gaussian (a) and a vectorial (b) PSF (σ ≈ 126 nm; emitter at the center of 31×31 bins of 25 nm; 4,000 ROIs per point; no background). Methods as in Fig. 1e,f.

### Drift correction and pupil analysis of the measured data

**Figure S8.**
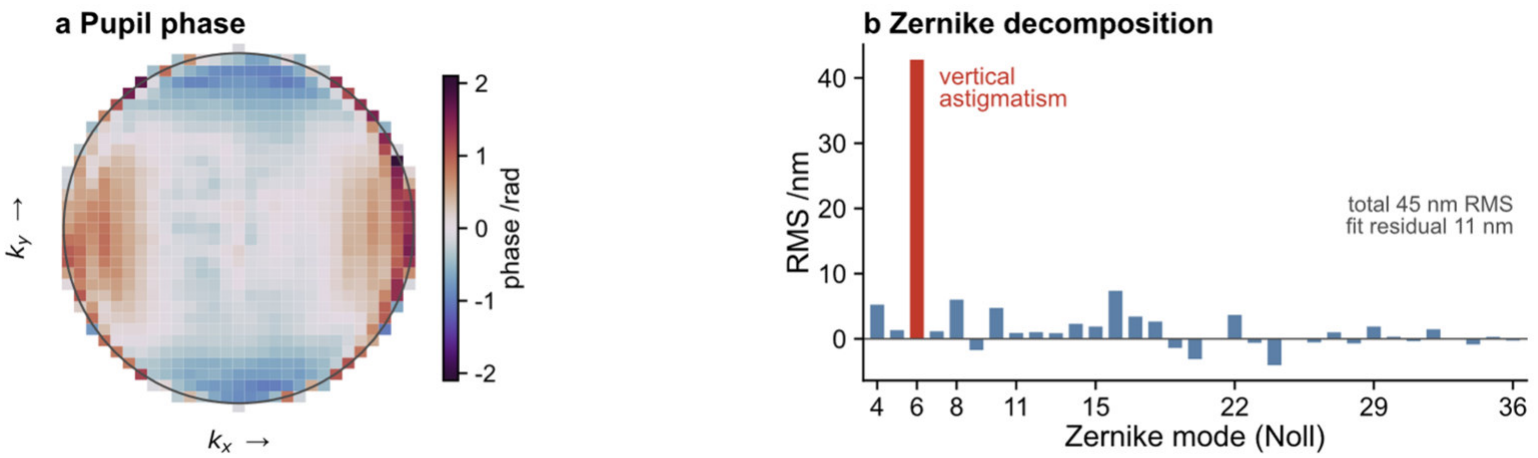
Calibrated PSF of the measured 3D data. a, Pupil phase from phase-only pupil retrieval (NA 1.49, λ = 668 nm; raw 55×55 pupil pixels). b, Its Zernike decomposition (Noll 4–36): vertical astigmatism dominates (43 of 45 nm RMS). This PSF is used for all reconstructions in Fig. 2.

**Figure S9.**
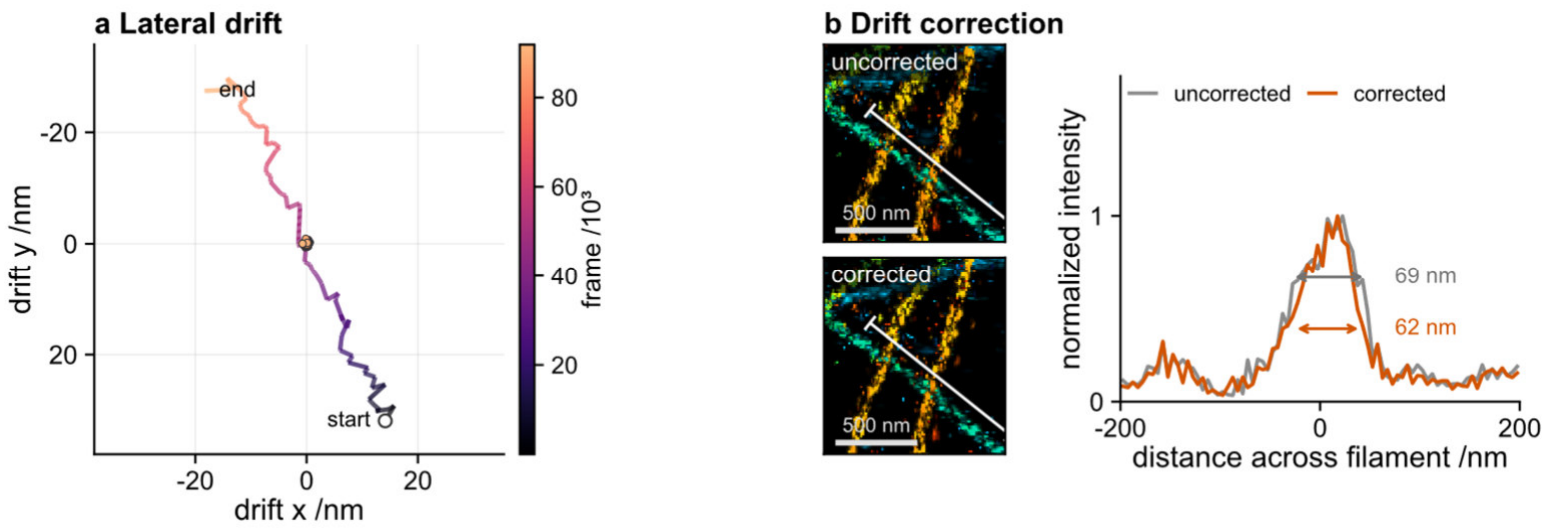
Lateral drift correction of the measured 3D data. a, Drift from cross-correlation of 60 MLLM block reconstructions (points) and the per-frame correction (line, colored by frame); 34 nm in x and 62 nm in y. Axial drift is not corrected. b, Region of Fig. 2c without and with the correction, and the profile across the marked filament (FWHM 69 and 62 nm). Scale bars, 500 nm.

### Equivalence of one-hot MLLM rendering to cSpline localization

Here we demonstrate the close conceptual agreement of a maximum likelihood fit, approximating the PSF by cSpline interpolation and assigning the coordinate to the nearest bin during rendering (histogramming) with picking the maximum voxel for each molecular detection in our MLLM map. Figure S10 compares reconstruction of these two strategies. It is seen that the results are virtually identical. Yet it should be noted that MLLM is non-iterative, whereas the fitting-based approach requires iteratively maximizing the likelihood (or minimizing the loss, being the negative log-likelihood).

**Figure S10.**
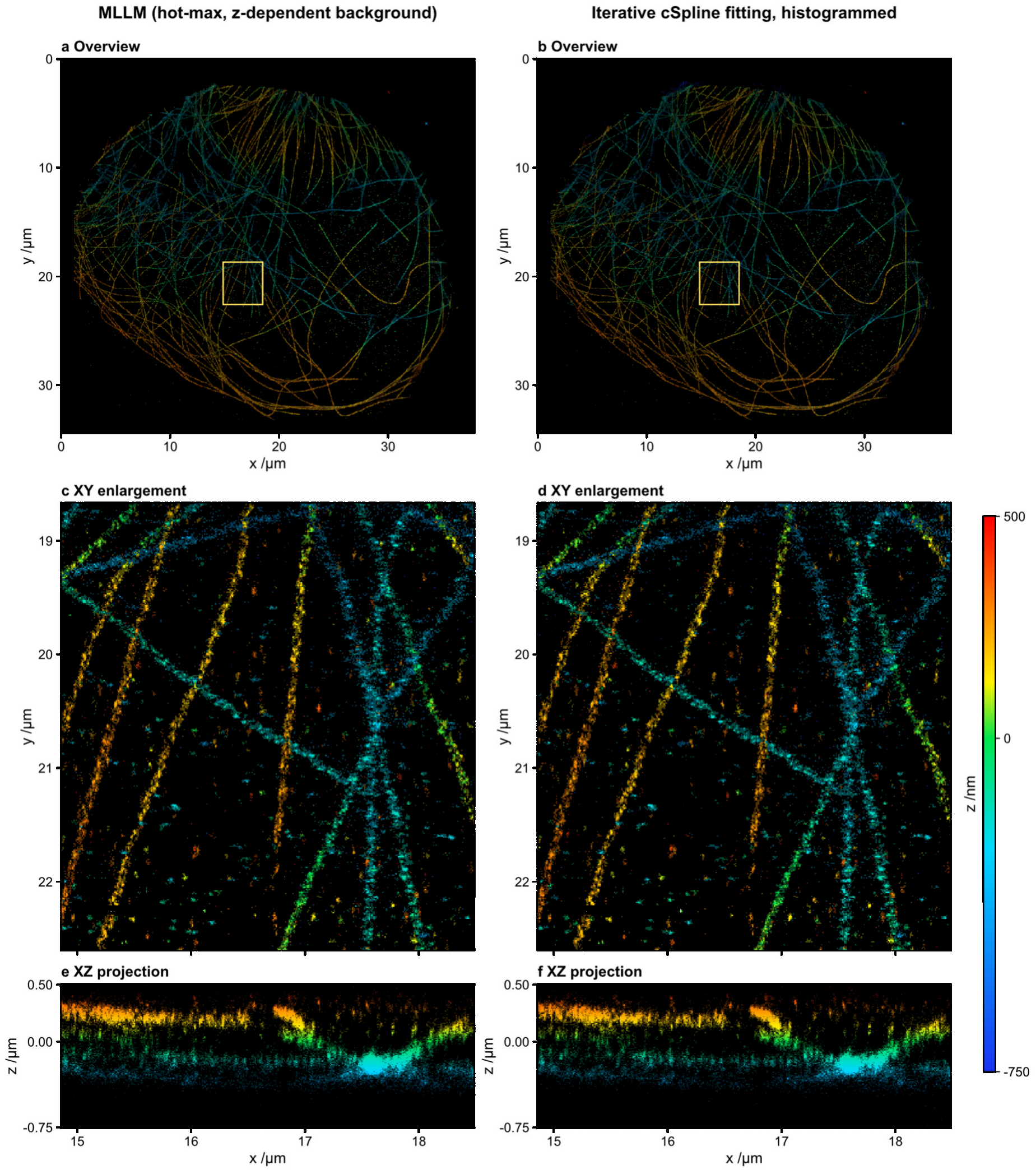
Drift-corrected measured experimental data 3D reconstruction with MLLM and iterative cSpline fitting. a,c,e, MLLM (non-iterative). b,d,f, Iterative cSpline fitting, histogrammed. Both methods use the same 3,572,756 retained 21 × 21-pixel ROIs from 91,946 available frames of the measured Tubulin-A647-3D dataset of the SMLM Challenge^8^ and a calibrated astigmatic PSF. MLLM places one count at each ROI’s joint 3D likelihood maximum; cSpline histograms coordinates, fitted with the authors’ original MATLAB/GPU software. Lateral drift (34 nm in x and 62 nm in y over the acquisition) was estimated by redundant cross-correlation of MLLM reconstructions of 60 consecutive ∼1,532-frame blocks and corrected identically for both methods: each MLLM likelihood map is shifted by a Fourier phase ramp before upsampling, and each cSpline coordinate by the drift of its frame; axial drift is not corrected. a,b, Full XY depth-color views. c,d, Enlargements of the boxed regions. e,f, XZ projections summed over the corresponding cropped y interval. Hue indicates the depth of maximum histogram count relative to the PSF center; the vertical colorbar spans −750 to +500 nm. All 151 planes (−750 to +750 nm) are retained numerically. Brightness is linear, with a full-field 97th-percentile cap per method for XY and a shared density cap for XZ. Sampling is 10.64 nm× 11.55 nm× 10 nm; coarse detector pixels are 117 nm× 127 nm. Scale bars: 5 µm (a,b) and 1 µm (c–f).

